# Comparison of HDO production from [2,3,4,6,6-^2^H_5_]Glucose and [^2^H_7_]Glucose as a marker of Glucose metabolism

**DOI:** 10.64898/2026.04.03.716329

**Authors:** Gaurav Sharma, Vinay Malut, Manoj Madheswaran, Haley Peters, Sameer Naik, Anna R. Nulk, Vikram D. Kodibagkar, James A. Bankson, Matthew E. Merritt

## Abstract

**PURPOSE:** Glycolytic production of HDO from the metabolism of perdeuterated glucose provides a means for metabolic imaging with ^2^H MRI. The present study compared HDO production from a cost-efficient [2,3,4,6,6-^2^H_5_]glucose with [^2^H_7_]glucose *in vitro* and *in vivo*.

**METHODS:** ^2^H NMR spectroscopy was performed to measure glucose consumption, lactate, and HDO production in the SFxL glioblastoma cell line. *In vivo* studies in healthy mice using ^2^H magnetic resonance spectroscopy were performed at 11.1 T after administering a bolus of either metabolic contrast agent. *In vivo* metabolite levels were quantified using unlocalized and slice-selective localized spectra.

**RESULTS:** Our *in vitro* results demonstrated similar glucose consumption and HDO production kinetics, although significant differences in lactate labeling were observed. The *in vivo* study showed comparable glucose consumption and HDO production kinetics following tail-vein bolus administration of either metabolic contrast agent, while lactate was not detected in the brain.

**CONCLUSION:** [2,3,4,6,6-^2^H_5_]glucose shows comparable HDO production to [^2^H_7_]glucose, while offering lower cost and reduced spectral complexity. These findings place [2,3,4,6,6-^2^H_5_]glucose as an alternative to [^2^H_7_]glucose for HDO-based DMI studies.

## 1 INTRODUCTION

Metabolic imaging provides direct insight into the metabolism of living systems in a minimally-invasive manner. The ability to visualize and quantify biochemical processes offers unprecedented insights into physiological function and pathological states such as cancer, diabetes, steatotic liver disease, cardiovascular disease, and neurological disorders. This technique links metabolic fluxes to cellular activity *in vivo*, addressing key questions in systems biology and medicine.

Numerous methods have been developed for metabolic imaging, each with its own advantages and drawbacks. Positron Emission Tomography (PET) is a gold-standard technique in clinical and preclinical settings. It leverages radiolabeled tracers to detect the uptake of a host of different substrates and antibodies. PET imaging with [^18^F]fluorodeoxyglucose (FDG-PET) can sensitively detect enhanced glucose uptake, a quality leveraged to produce a ubiquitous tool for cancer detection, staging, and monitoring. However, FDG-PET imaging uses ionizing radiation and cannot detect downstream metabolism of the tracer. Magnetic Resonance Spectroscopy (MRS) can offer similar information to PET without the use of ionizing radiation, albeit with lower sensitivity. ^13^C-MRS is a widely adopted technique in clinical research and preclinical settings, providing dynamic metabolic information using ^13^C-labeled substrates. ^13^C-MRS exhibits low sensitivity (low signal-to-noise ratio, SNR), resulting in long scan times thereby limiting its practicality for clinical use. The use of hyperpolarized ^13^C-MRS circumvents this bottleneck by boosting the SNR by 10-40,000 times, resulting in rapid data acquisition and real-time metabolic imaging. Despite the benefits of HP-^13^C MRS, it is limited by the short T_1_ of the substrates, the complex delivery of the metabolic contrast agent, and the need for a separate prepolarizer for sample preparation.

Recently, deuterium metabolic imaging (DMI) emerged as a competitive metabolic imaging technology. It utilizes exogenous, non-radioactive, deuterium-labeled metabolic contrast agents, and provides information on both uptake and downstream metabolism. The availability of diverse deuterium-labeled compounds makes it an attractive imaging modality to study multiple metabolic pathways. Although the deuterium nucleus has a low gyromagnetic ratio (□1/6 of ^1^H) and therefore low relative sensitivity, this can be compensated for by the ability to acquire more signal averages in a given imaging paradigm due to the short T_1_ endemic to quadrupolar nuclei (spin > ½). Low natural abundance (0.0015%) brings another advantage in the form of low background signal from endogenous metabolites. [6,6-^2^H_2_]glucose is a widely used agent employed in various animal and human studies, but even with the fast repetition time for deuterium experiments the agent is limited in functionality by long acquisition times.

This issue can be addressed in two ways. First, advancements in DMI hardware, such as the use of a higher magnetic field, designing different geometries of deuterium coils, and altering the acquisition trajectories for imaging and post-processing. A second option to boost SNR is to use a higher number of deuterium labels per glucose molecule. Mahar et. al. reported the use of [^2^H_7_]glucose (uniform deuterium enrichment) to study the metabolism of the healthy rat brain via production of deuterated water (HDO). This study suggested that region-specific HDO signal could be used as a metabolic biomarker reflecting not only glucose uptake, as observed with FDG-PET, but also glucose metabolism. In another study, Cocking et. al. performed a comparison between [^2^H7</SUP>]glucose and [6,6- ^2^H2]glucose in human participants. [^2^ H7]glucose metabolism resulted in a significant increase in HDO, Glx (Glutamate and Glutamine), and lactate signal compared to [6,6- ^2^H2]glucose. These studies demonstrate an advantage to using [^2^H7]glucose over [6,6- ^2^H2]glucose. However, [^2^H7]glucose costs 10-15x more than [6,6- ^2^H2]glucose, presenting a barrier to clinical use.

Recently, Zou et. al. reported the synthesis of a new tracer, [2,3,4,6,6- ^2^H5]glucose (hereafter [^2^H5]glucose), using a synthetic route that is 10x cheaper than the synthesis of [6,6- ^2^H2]glucose. The additional deuterium labelling at C2, C3, and C4 positions offered a 2.41x SNR increase in glucose signal compared to [6,6- ^2^H2]glucose in a model of rat glioma. The additional deuterium labeling also led to higher production of HDO through glycolysis. This tracer is both more affordable and liberates more HDO compared to [6,6- ^2^H2]glucose, offering an accessible route to selectively resolve the glycolytic pathway.

No study has yet reported a direct comparison between [^2^H5]glucose and [^2^H7]glucose *in vivo*. The affordability of [^2^H5]glucose along with high production of HDO may accelerate DMI as a clinical imaging paradigm. This gap in the existing literature prompted us to conduct a systematic evaluation of contrast agent performance in both *in vitro* and *in vivo* models.

## 2 METHODS

### 2.1 CHEMICALS AND REAGENTS

[2,3,4,6,6- ^2^H5]glucose (Lot No. DLM-11743-PK) was received as a gift from Cambridge Isotopes Laboratory (CIL), and [^2^H7]glucose (DLM-2062-5) was purchased from CIL. Dulbecco’s Modified Eagle Medium (DMEM), Dimethyl Sulfoxide (DMSO), and Phosphate Buffered Saline (PBS) were purchased from Thermo Scientific (Waltham, MA, USA). Fetal Bovine Serum (FBS) was purchased from Atlas Biological (Fort Collins, CO, USA). Deuterated pyrazine (Pyrazine-d4) and deuterium oxide (D2O) were purchased from Sigma Aldrich (St. Louis, MO, USA). Isoflurane anesthesia was purchased from Patterson Veterinary (Ocala, FL, USA). The deuterated glucose were prepared by dissolving in sterile-filtered saline solution. A 0.9% sterile-filtered saline solution was prepared from NaCl purchased from Thermo Fisher Scientific (Waltham, MA, USA), and, for *in vivo* studies, was constituted with 1% v/v sterile heparin (50 USP/mL, final concentration) to prevent clotting. All reagents were sterile-filtered before administration to cells or animals.

### 2.2 CELL CULTURE AND STABLE ISOTOPIC TRACING

The glioblastoma SFxL cell line used in this study was generously gifted by Dr. Ralph Deberardinis from the University of Texas Southwestern Medical Center. SFxL cells were maintained in Dulbecco’s modified Eagle medium (DMEM) supplemented with penicillin–streptomycin (50 μg/mL each) and 10% v/v FBS at 37°C in a humidified atmosphere containing 5% CO2 in an incubator (Heracell Vios 160i). Media was replaced daily, and cells were split (1:10) at 70-80% confluency. At passage 6, cell counting was performed, and (5 × 10^6^) cells were seeded in 10 cm^2^ culture plates for the tracer study. Before introducing the tracers, the cells were maintained in their respective media until 70-80% confluency was achieved (∼16 hr). After this period, standard media were replaced with self-made media of the same composition, in which unlabeled glucose was replaced with either [^2^H7]glucose or [^2^H5]glucose tracers (6 mL of 8.5 mM concentration) for six hours.

During this incubation period, media samples (200 μL aliquots) were collected at 30 min, 60 min, 120 min (2 hr), 180 min (3 hr), 360 min (6 hr), and stored at −80 °C for further analysis. At the end of the 6 hr incubation, all media were collected. Cells were washed with warm PBS and harvested by trypsinization. Cell counting was performed on 20 μl of the resulting suspension. The remaining samples were centrifuged for 5 minutes at 300 × g at 4 °C, followed by aspiration of the supernatant. The cells were then resuspended in 4 mL of saline, centrifuged for 5 min at 300 × g, and followed by a final aspiration of the supernatant. The remaining washed cell pellet was immediately immersed in liquid nitrogen to preserve metabolic profiles for ^2^H nuclear magnetic resonance analysis.

### 2.3 NMR STUDY

Media samples were analyzed directly without performing metabolite extraction. The NMR sample is prepared by mixing 45 μL of the media samples and 5 μL of the 25 mM pyrazine-d4 internal standard stock solution to attain a 50 μL total sample volume and a final concentration of 2.5 mM pyrazine-d4. The addition of pyrazine-d4 internal standard permitted us to quantify HDO, ^2^H-lactate, unconsumed [^2^H5]glucose, and [^2^H7]glucose from ^2^H-NMR data. Finally, the sample was transferred into 1.7 mm NMR sample tubes for NMR data acquisition.

NMR experiments were executed at 14 T (600 MHz) (Bruker Biospin with Neo console) using a 1.7 mm TCI Micro CryoProbe. ^2^H NMR experiments were acquired using the following acquisition parameters: spectral width 2000 Hz, flip angle 90°, TD 3072, D1 1 s, and total acquisition time 1.5 hr.

### 2.4 ANIMAL STUDY

All animal studies were reviewed and approved by our Institutional Animal Care and Use Committee which is accredited by AAALAC International. Eight week old C57Bl6/J mice were purchased from the Jackson laboratory (USA) and acclimatized in the McKnight Brain Institute animal facility one week prior to imaging. Mice were fed standard chow *ad libitum*. Mice were anesthetized initially with 2% isoflurane mixed with oxygen gas and then maintained at 1.2-1.5%. For administering the bolus of metabolic contrast agents, the tail vein was catheterized using a 30-gauge needle. The body temperature of the mice was maintained at 37 ºC using a heated circulating water pad. The respiratory pad was placed close to the mouse’s heart to monitor the respiration rate during the entire period of the experiment.

### 2.5 MAGNETIC RESONANCE IMAGING SYSTEM AND SCAN DETAILS

All MR studies were executed with an 11.1 T magnet (Magnex Scientific, Yarnton, United Kingdom) connected to a Bruker Avance III HD controlled via ParaVision 6.0.1 software (Bruker BioSpin, Billerica, MA). The system was equipped with an RRI BFG-240/120-S6 gradient coil, featuring a 120-mm bore size, a gradient strength of 100 mT/m, and a rise time of 200 μs. To ensure a homogeneous magnetic field around the mouse brain, shimming was optimized by acquiring a B0 field map, and if needed, a second-order shim correction was performed using mapshim with an ellipsoid region that includes mainly the brain, using a custom-built 85-mm, actively decoupled, linear-volume ^1^H transmit-receive coil tuned to the ^1^H-NMR frequency of 470 MHz. A custom-built 14-mm diameter circular transmit-receive surface coil resonating at 72.26 MHz was utilized to acquire ^2^H-MRS/I. 90-degree pulse calibration was done prior to the animal study using 2% D2O phantom. The MR acquisition parameters are provided in Table 1.

**Table 1:** T2 Rare spin echo ^1^H MRI and ^2^H MRS acquisition parameters.

| Acquisition Parameters | Anatomical $^1\text{H}$ MRI | Non-localized $^2\text{H}$ MRS | Slice-selective localized $^2\text{H}$ MRS |
| --- | --- | --- | --- |
| Field of View (FOV) | 32 x 32 mm<sup>2</sup> | - | - |
| Matrix Size | 128 x 128 | - | - |
| Slice Thickness | 2 mm | - | 5.5 mm (Coronal) |
| Number of Slices | 8 | - | 1 |
| Repetition Time (TR) | 3000 ms | 300 ms | 100 ms |
| Echo Time (TE) | 40.5 ms | - | 1.05 ms |
| Flip Angle (FA) | $90^\circ$ | $60^\circ$ | $60^\circ$ |
| Rare Factor | 16 | - | - |
| Number of Averages | 4 | 256 | 1024 |
| Bandwidth | - | 1000 Hz | 3000 Hz |
| Number of Samples | - | 256 | 512 |
| Scan Duration | 3 min 12 s | 1 min 16 s | 1 min 56 s |

### 2.5 ^2^H NMR AND ^2^H MRS PROCESSING

MestReNova v14.0.1-23284 (Mnova, Mestrelab Research S.L.) was used to post-process the ^2^H NMR and ^2^H MRS spectra. After importing the FID into Mnova, apodization, zero filling, phase correction, and baseline correction were performed in batch to ensure consistent processing across the same sample. After post-processing, line fitting was performed using a Lorentzian-Gaussian peak fitting method. For *in vitro* studies, concentrations of HDO, ^2^H-lactate, residual [^2^H5]glucose, and residual [^2^H_7_]glucose i the cell media were calculated relative to peak areas of the internal standard pyrazine-d_4_ (2.5 mM). For *in vivo* studies, the concentrations of metabolites were calculated using the background HDO signal, assuming 80% water in mouse brain tissue corresponds to 13.2 mM HDO.

### 2.6 STATISTICS

All statistical analysis for this research were conducted using GraphPad Prism (La Jolla, CA, USA, version 10.1.2). Statistical significance was established by using an unpaired two-tailed t-test with Welch correction and no correction for multiple comparisons. Data are represented as mean ± SD (n = 3).

## 3 RESULTS

### 3.1 *IN VITRO* comparison of [^2^H_5_]glucose and [^2^H_7_]glucose

Media samples collected at different time points, from SFxL cells treated with [^2^H_5_]glucose and [^2^H_7_]glucose, were analyzed by ^2^H NMR. During the course of study, downstream metabolism of [^2^H_5_]glucose and [^2^H_7_]glucose resulted in the appearance of lactate and HDO (shown in Figures 1A and B, respectively).

**FIGURE 1:**
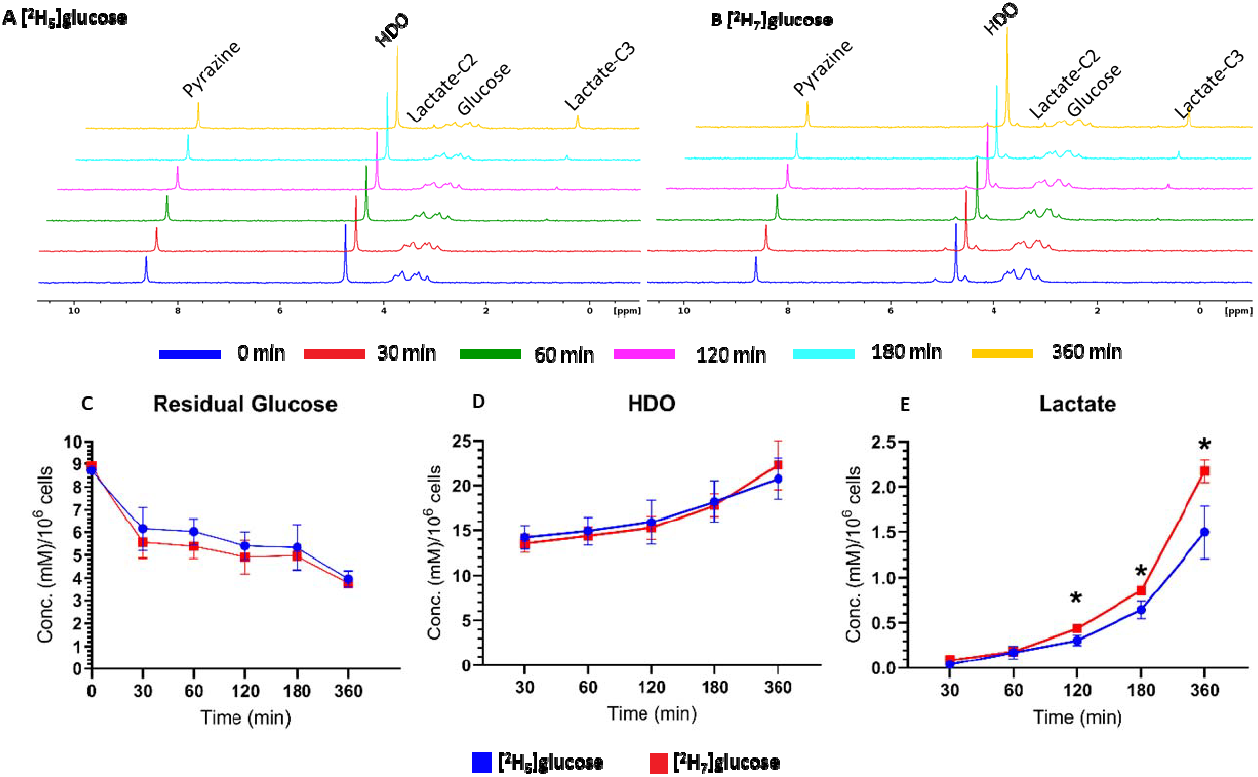
Stacked ^2^H NMR Spectra of Culture Media Collected at 0 min (BLUE), 30 min (RED), 60 min (GREEN), 120 min(PINK), 180 min (CYAN), and 360 min (DARK YELLOW) Minutes following incubation of SFxL Glioblastoma cells with (A) [^2^H_5_]glucose (B) [^2^H_7_]glucose. A 0.1-ppm horizontal shift was applied to each spectrum to improve clarity and permit visual comparison of temporal metabolic changes between the two agents. Quantitative analysis of metabolite concentrations using pyrazine-d_4_ as an internal standard over time for the two tracers. **(C)** Glucose concentrations steadily declined across all time points with no significant difference between [^2^H_7_]glucose (red) and [^2^H_5_]glucose (blue). **(D)** HDO production increased continuously throughout the incubation period, reflecting ongoing glycolytic and oxidative activity under both tracer conditions. **(E)** Labeled lactate concentrations rose progressively over time, with [^2^H_7_]glucose showing a more pronounced increase at later time points compared with [^2^H_5_]glucose. Data are presented as the means ± SD of N = 3 biological replicates. Statistical significance was established by using an unpaired two-tailed Welch’s t-test. No multiple-comparison correction was applied. Significance is \**P* ≤ 0.05.

Glucose consumption did not differ between [^2^H_5_]glucose and [^2^H_7_]glucose as indicated by no change in ^2^H NMR signal of residual glucose in media samples (Figure 1C). Interestingly, the production of HDO is also unchanged between the two tracers at all time points, while [^2^H_7_]glucose did trend higher at 360 min (Figure 1D). Incorporation of [^2^H_7_]glucose into labeled lactate increased over time and is significantly enriched from 120 to 360 minutes compared to [^2^H_5_]glucose (Figure 1E).

### 3.2 *in vivo* comparison of [^2^H_5_]glucose and [^2^H_7_]glucose

For the *in vivo* study, both deuterated glucose agents were administered via tail vein injection. Non-localized and localized ^2^H MRS experiments were acquired prior to administration to assess the natural abundance background HDO signal for normalization. After bolus injection, a series of non-localized spectroscopy scans (□1.3 minutes/spectrum) were performed to track the appearance of agents in the brain. For ^2^H localized spectroscopy, the coronal slice was adjusted to cover the whole brain, carefully avoiding the phantom (Figure 2A). Typical spectra for the uptake and consumption of the agents demonstrate the delivery and metabolism of the agents to produce HDO (Figure 2B,C). By spectral deconvolution, we estimated the amount of deuterated agent consumed and production of HDO resulting from metabolic utilization (Figure 2D). Trends observed in cell culture mirror *in vivo* results. Total glucose consumption did not significantly differ (*P* > 0.05) between [^2^H_5_]glucose and [^2^H_7_]glucose at most of the time points as indicated by residual glucose signal (Figure 2E). HDO production was also unchanged though it trended higher with [^2^H_7_]glucose (Figure 2F).

**Figure 2:**
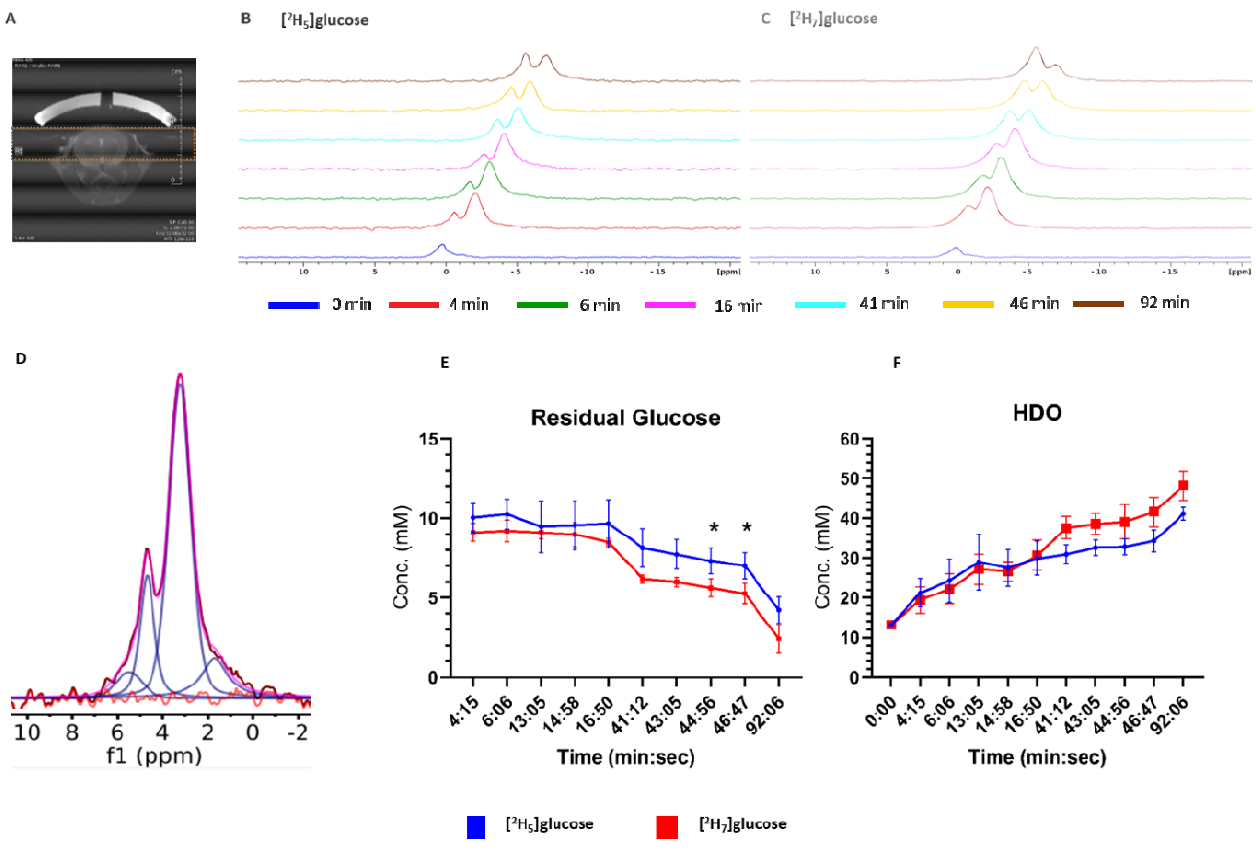
(A) Anatomical T_2_-weighted MRI demonstrating the slice-selection geometry used to encompass the entire brain for ^2^H slice-selective spectroscopy. (B) Time-resolved, localized *in vivo* ^2^H MR spectra acquired from the mouse brain, using a ^2^H transmit-receive surface coil, following bolus injection of ^2^H_5_-glucose and (C) ^2^H_7_-glucose. ^2^H spectra acquired at successive pre-injection (blue) and post-injection time points (4 min (red), 6 min (green), 16 min (pink), 41 min (cyan), 46 min (yellow), 92 min (brown). The gradual decrease of glucose resonances and concurrent changes in downstream deuterated metabolite signals (HDO) reflect glucose consumption. (D) Representative line-fitted slice-selective ^2^H spectrum using a multi-peak line-shape model to quantify the HDO (□4.7 ppm) and glucose (□3–4 ppm) resonances [individual fitted peak (navy blue), sum (magenta), residual (red)]. (E) Residual glucose concentrations steadily declined across most of the time points. (F) HDO production increased continuously throughout the imaging period, reflecting ongoing glycolytic and oxidative activity under both agent conditions. Data are presented as the means ± SD of n = 3 biological replicates. Statistical significance was established by using an unpaired two-tailed Welch’s t-test. No multiple-comparison correction was applied. Significance is \**P* ≤ 0.05.

## 4 Discussion and Conclusions

In this study, we compared [^2^H_5_]glucose and [^2^H_7_]glucose as metabolic contrast agents in both *in vitro* and *in vivo* models. First, we evaluated these agents in a cell culture system using human glioblastoma SFxL cells. The ease of handling and controlled experimental environment make cell culture a suitable platform for initial tracer evaluation due to reduced biological complexity and lower variability. Additionally, SFxL cells represent a highly glycolytic cancer model with an increased demand for glucose to support rapid proliferation and metabolic activity.

Our cell culture results demonstrate comparable uptake of the two tracers (*P* > 0.05) at almost every time point. The glucose consumption also resulted in similar levels of HDO production despite the loss of two deuterium labels for [^2^H_5_]glucose (Figure 1D). In contrast, significant differences (*P* < 0.05) were seen in lactate signal between the tracers at time points 120, 180 and 360 minutes, which can be attributed to the additional deuterium labeling in [^2^H_7_]glucose at the C1 position contributing to the lactate C3 position. These findings indicate that the new tracer performs comparably to [^2^H_7_]glucose when HDO is used as the primary metabolic biomarker. The [^2^H_7_]glucose provides higher SNR ratio for lactate C3 as it results in the production of an additional single deuterium at this position derived from the C1 position. If lactate labeling at the C3 position is the target for investigation, [^2^H_7_]glucose may be the superior agent from a sensitivity standpoint. The insignificant difference in HDO production between the two agents is explained by the appearance of deuterium in the C3 position of lactate. Our data demonstrates that HDO production primarily arises from the reactions in glycolysis prior to lactate production. The persistence of methyl labeling at long time points demonstrates that lactate functions as a “trap” for the labeling derived from the C1 and C6 positions. Deuterium derived from positions H2-H5 is readily eliminated as HDO by glycolytic flux.

To assess whether these trends translate *in vivo*, we evaluated the metabolism of both tracers in the brains of healthy eight-week-old C57BL/6J mice. Given the brain’s high glucose demand, this system provides a sensitive model for evaluating tracer uptake and metabolic turnover. Consistent with cell culture results, glucose uptake in the brain was comparable between the two agents. Importantly, the downstream metabolic product HDO remained similar between tracers across all time points (*P* > 0.05). Our *in vitro* and *in vivo* results indicate that the use of [^2^H_7_]glucose can be replaced with [^2^H_5_]glucose when measuring HDO production as the principal metabolic readout. We did not observe ^2^H-lactate or ^2^H-Glx (glutamate/glutamine) signals derived from either tracer even at late time points. Possibly this is due to the smaller mouse brain volume compared with the rat brain, which results in lower SNR for these metabolites. ^2^H-Glx (glutamate/glutamine) can be masked with glucose signal *in vivo* because of the broad linewidth of ^2^H MR spectra and may result in overestimation of residual glucose.

In addition to its improved cost profile and comparable metabolic handling, [^2^H_5_]glucose also offers advantages in spectroscopic analysis. The absence of C1 labelling allows for straightforward quantification of HDO due to the lack of anomeric glucose signal, which ordinarily resonates near the HDO signal and is difficult to deconvolute *in vivo*.

Deuterium metabolic imaging (DMI) is emerging as a competitive metabolic imaging technology and holds promising potential for translation into clinical settings. SNR challenges with ^2^H detection have prevented detection of signal at high spatial resolutions, as more tissue is needed to generate sufficient metabolic product for detection. The HDO metabolic pool is large and changes are easily detectable. The use of [^2^H_7_]glucose maximizes the HDO production reported in previous studies. The present study demonstrates that [^2^H_5_]glucose offers a more cost-effective alternative to [^2^H_7_]glucose, without sacrificing SNR for HDO production. The absence of C1 and C5 labeling allows for more accurate HDO quantification, as it reduces spectral complexity. If experiments are executed in such a manner (imaging immediately after glucose injection), [^2^H_5_]glucose should provide an inexpensive alternative for imaging glycolytic flux by DMI.

## Acknowledgement

A portion of this work was performed at the National High Magnetic Field Laboratory, which is supported by National Science Foundation Cooperative Agreement number DMR-1644779, & the State of Florida.

## Author contributions

**Conceptualization:** GS, VM, MEM; **Methodology**: GS, VM; **Data Collection**: GS, VM, MM, HP; **Formal Analysis**: GS, VM, MM, HP, SN, AR, MEM **Writing—original draft**: GS; **Writing—review & editing**: GS, VM, MM, HP, SN, AR, VK, JB, MEM; **Supervision**: MEM; **Resources**: MEM; **Funding Acquisition**: MEM

